## Supplementary Information for "HAPP: High-Accuracy Pipeline for Processing deep metabarcoding data"

---

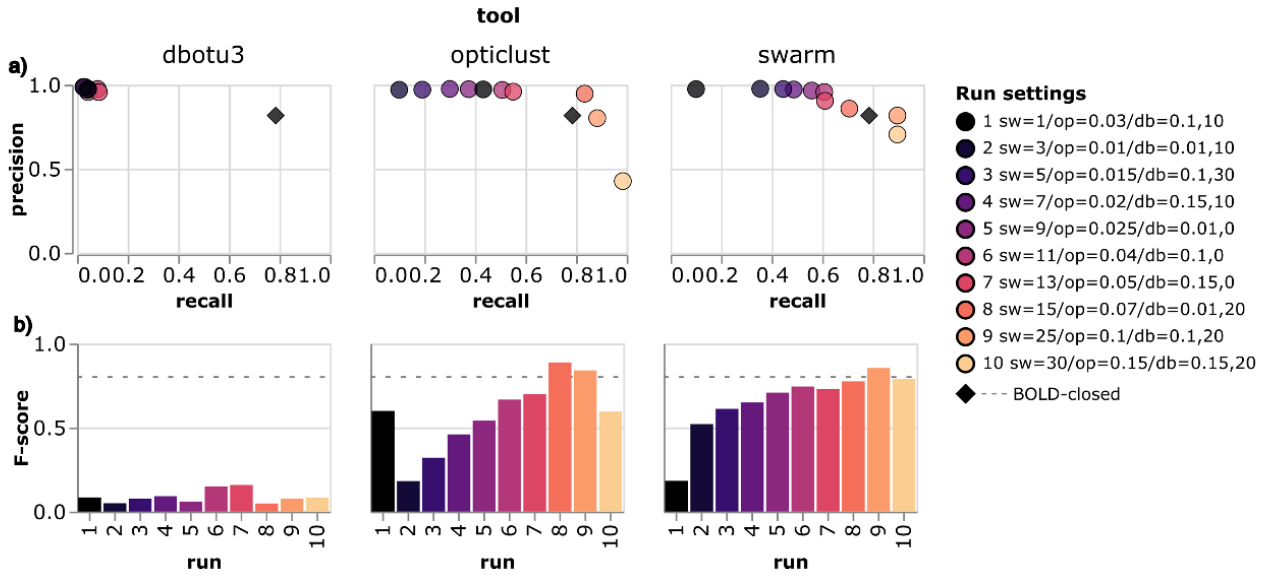


**Fig. S1**. Benchmarking of ASV clustering tools, using the original species-level annotation of the BOLD sequences (rather than the GBIF-resolved BIN-level annotations), and assessing the putative performance of the RESL algorithm on IBA data through closed reference clustering according to the SINTAX assignments to BOLD BINs (‘BOLD-closed’). **a**. Precision vs. recall for each run of the different tools. Colors correspond to the different runs and the legend summarizes the settings used: ‘sw’ gives the difference cutoff in Swarm, ‘op’ the distance cutoff in OptiClust, and ‘db’ the distance and abundance cutoffs in dbOTU3. The diamond symbol shows the result obtained in closed reference clustering using BOLD BIN assignments. **b**. F-score values (harmonic mean of precision and recall) of the different runs for each tool. The dashed line shows the F-score for closed reference clustering using BOLD BIN assignments.

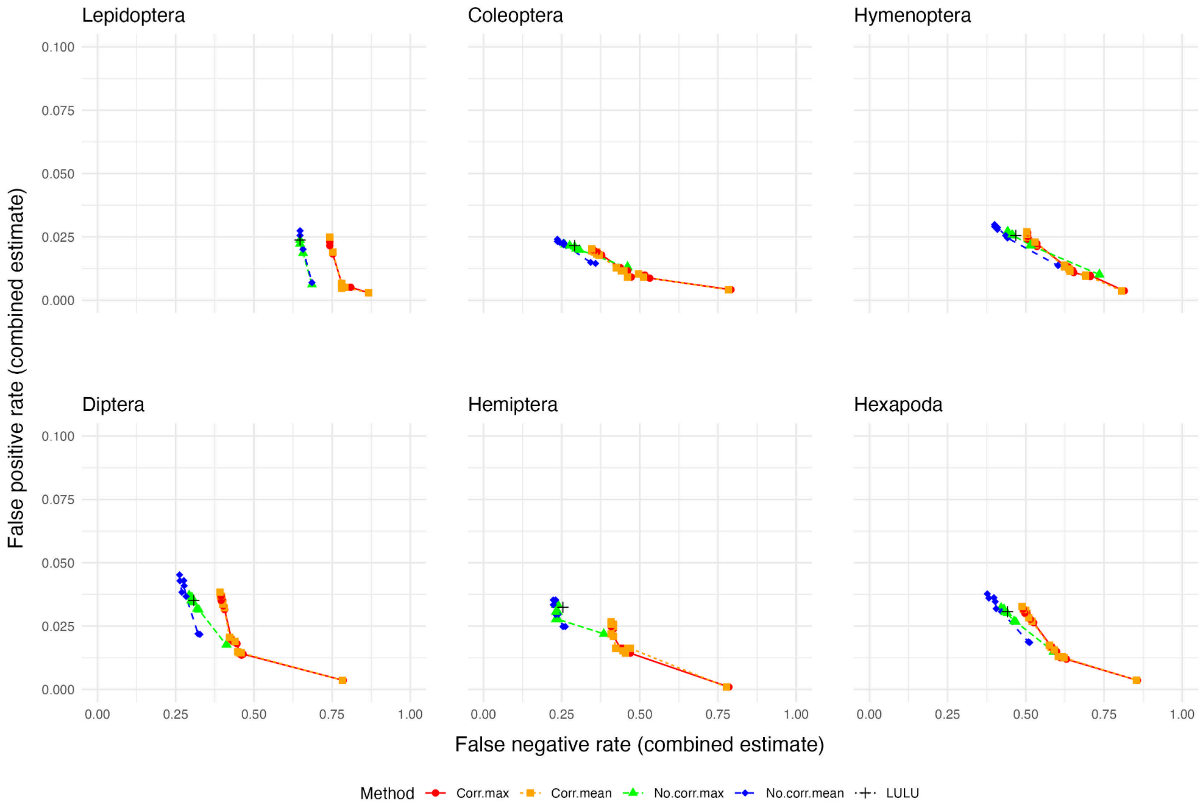


**Fig. S2. Performance of the echo algorithm of NEEAT**. The plots show the performance on various hexapod taxa in terms of our combined estimates of false positives (authentic CO1 sequences removed) and false negatives (false CO1 sequences remaining), and compares it to the performance of LULU. Four different settings were explored: 'Corr.max' = enforcing a significant correlation in read numbers between noise and signal, varying max read ratio from 0.05 to 1.0. 'Corr.mean' = ditto, varying mean read ratio from 0.05 to 1.0. 'No.corr.max' = no correlation test, varying max read ratio threshold from 0.1 to 1.0. 'No.corr.mean' = ditto, varying mean read ratio from 0.1 to 1.0. 'LULU' = LULU algorithm with default settings (similar to 'No.corr.max' with read ratio of 1.0).

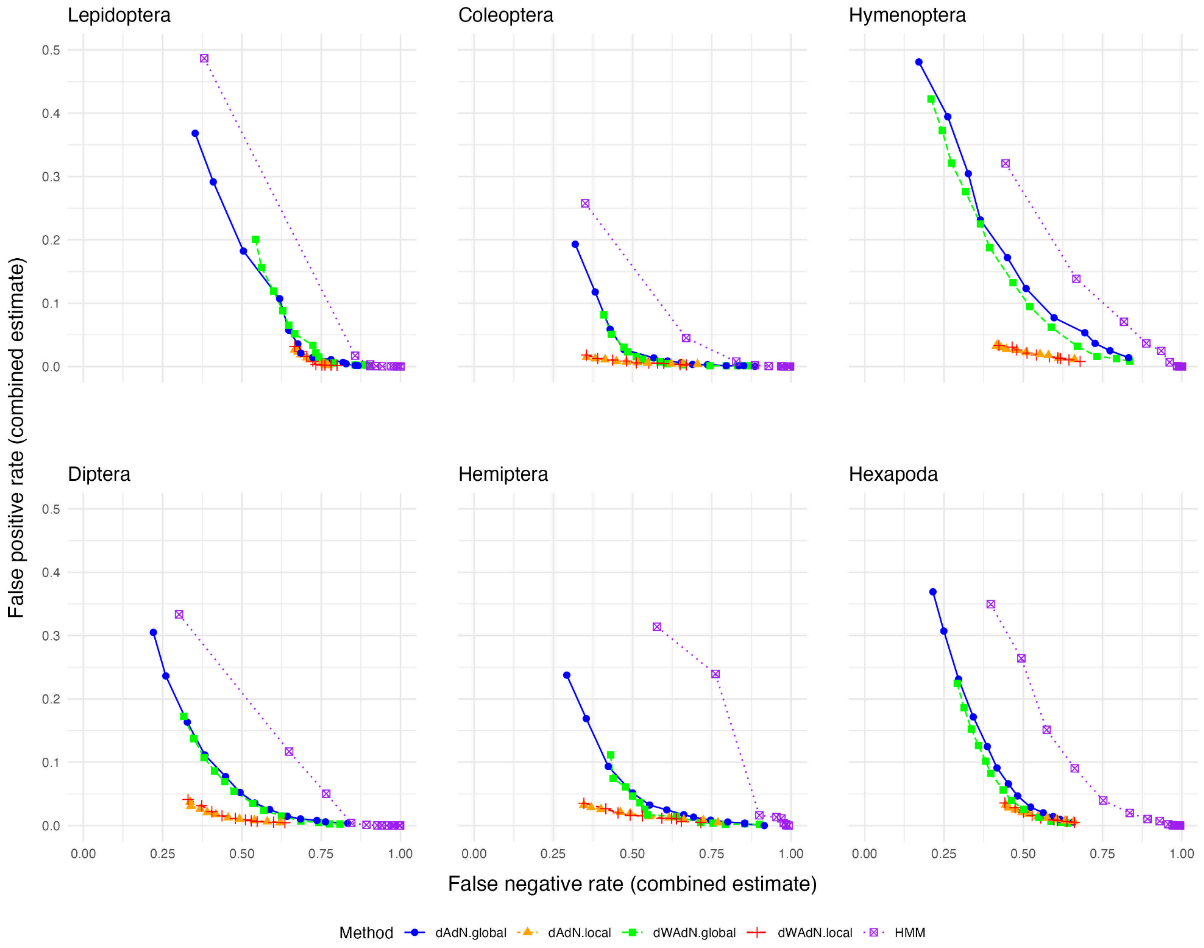


**Fig. S3. Performance of the evolutionary signature algorithm of NEEAT**. The plots show the performance on various hexapod taxa in terms of our combined estimates of false positives (authentic CO1 sequences removed) and false negatives (false CO1 sequences remaining), and compares it to the performance of the HMM profile algorithm. Four different settings were explored: ‘dAdN.global’ = unweighted amino acid distances, comparing sequences across all samples, varying distance threshold from 1.0 to 6.0. ‘dAdN.local’ = ditto, comparing sequences with sample overlap, varying distance threshold from 0.4 to 2.7. ‘dWAdN.global’ = biochemically weighted amino acid distances, comparing sequences across all samples, varying distance threshold from 0.8 to 4.0. ‘dWAdN.local’ = ditto, comparing sequences with sample overlap, varying distance threshold from 0.1 to 1.7. ‘HMM’ = HMM profile algorithm, varying bitscore threshold from 160 to 300.

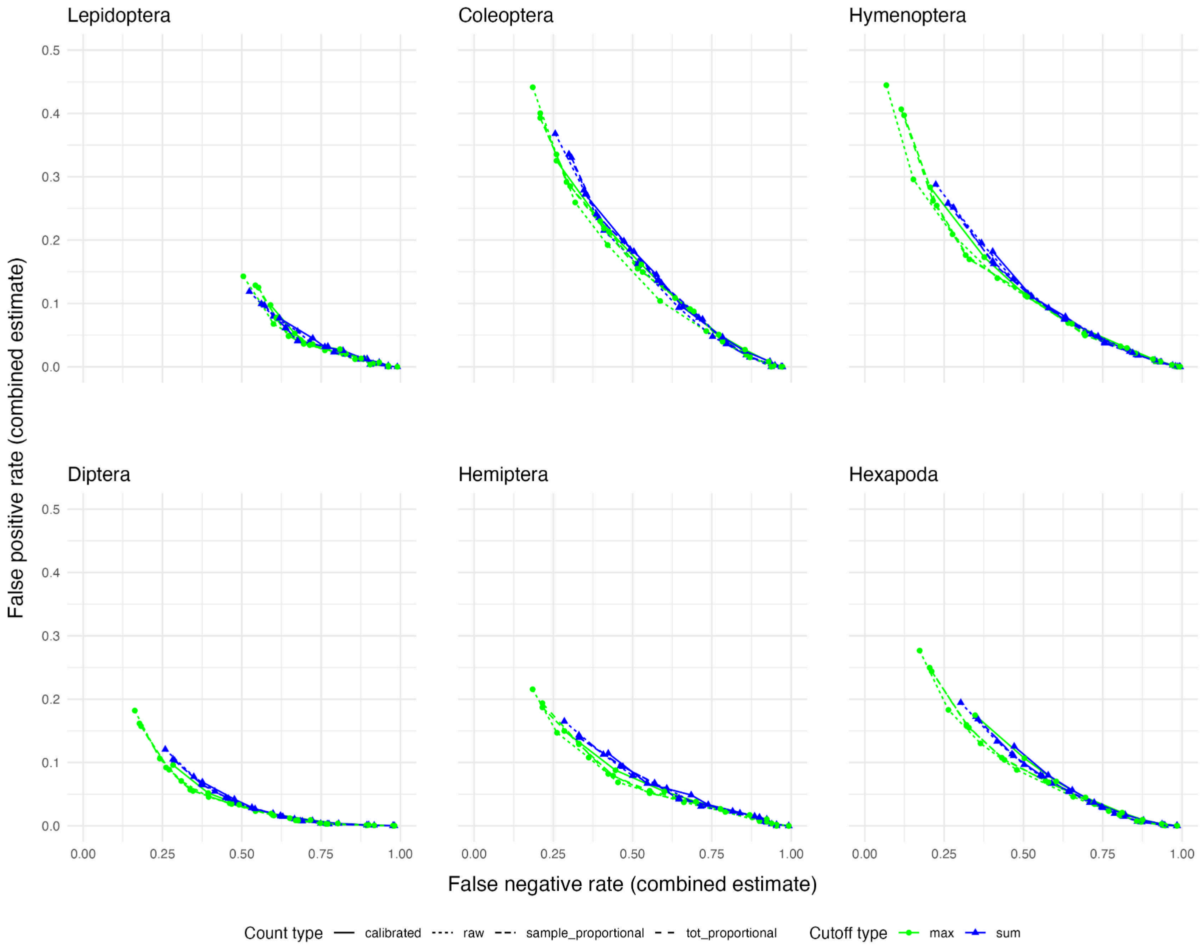


**Fig. S4. Performance of the abundance threshold algorithm of NEEAT**. The plots show the performance on various hexapod taxa in terms of our combined estimates of false positives (authentic CO1 sequences removed) and false negatives (false CO1 sequences remaining). We explored four different types of read counts: 'calibrated' = read counts calibrated by the number of biological spike in reads. 'raw' = raw read counts. 'sample_proportional' = the proportion of the reads in the sample, with the biological spike in reads excluded. 'tot_proportional' = the proportion of the reads in the sample, with biological spike in reads included. We explored two types of thresholds: 'max' = maximum reads across samples. 'sum' = the sum of reads across samples. The threshold was varied from 1 to 500; for proportional counts, the threshold was divided by 1E6 (the expected number of reads per sample).

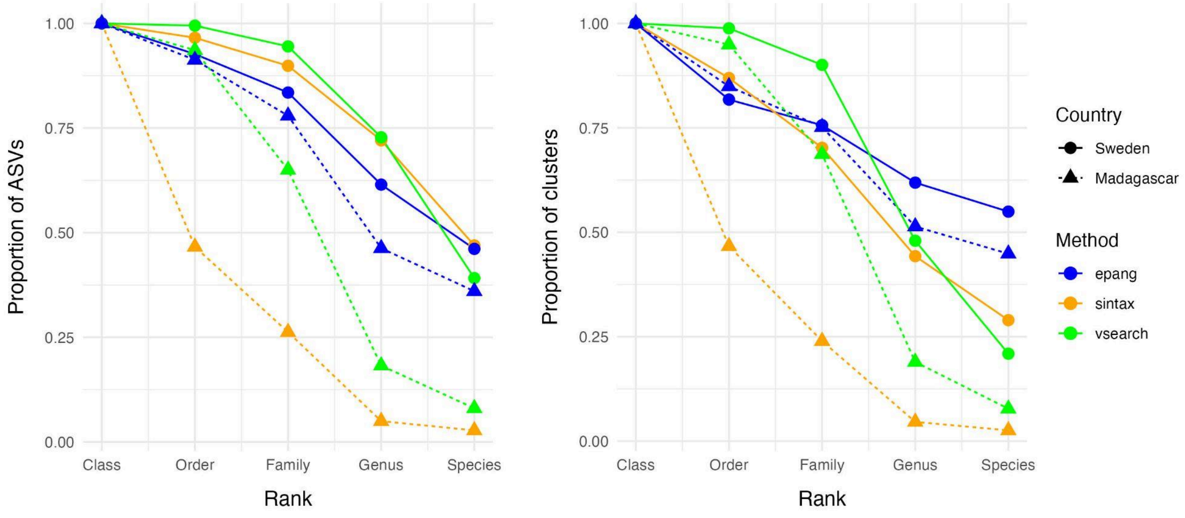


**Fig. S5. Taxonomic annotation success of different algorithms for faunas with different coverage in reference databases.** The plots show the annotation success for IBA data at taxonomic ranks ranging from phylum to species for a k-mer based method ('sintax'), an alignment-based method ('vsearch'), and a phylogenetic placement method ('epang'). Results are shown for a fauna well covered in the BOLD reference database and the reference tree used for annotation ('Sweden') and for a poorly covered fauna ('Madagascar'). The results for clusters are based on the taxonomic annotation for the representative ASV of each cluster.
